# Dysfunctional glymphatic system with disrupted aquaporin-4 expression pattern on astrocytes causes bacterial product accumulation in the CSF during pneumococcal meningitis

**DOI:** 10.1101/2022.04.14.488283

**Authors:** Jaqueline S. Generoso, Sigrun Thorsdottir, Allan Collodel, Diogo Dominguini, Roberta R. E. Santo, Fabricia Petronilho, Tatiana Barichello, Federico Iovino

## Abstract

**Background:** Pneumococcal meningitis, inflammation of the meninges due to an infection of the Central Nervous System caused by *Streptococcus pneumoniae* (the pneumococcus), is the most common form of community-acquired bacterial meningitis globally. The brain is separated from the systemic circulation by the blood-brain barrier (BBB), and meningitis triggers the host immune response increasing the BBB permeability, allowing peripheral immune cells to reach the cerebrospinal fluid (CSF), and increasing debris production. The glymphatic system is a glial-dependent clearance pathway that drives the exchange of compounds between the brain parenchyma and the CSF regulating the waste clearance away from the brain. Aquaporin-4 (AQP4)-water channels on astrocytic end feet regulate the solute transport of the glymphatic system.

**Methods:** Wistar rats, either subjected to pneumococcal meningitis or to artificial-CSF (sham), received Evans blue albumin (EBA) intracisternal. Overall, the meningitis group presented a significant impairment of the glymphatic system by retaining the EBA in the brain without consistently releasing the EBA into the bloodstream compared to the sham non-infected group. Through western blot and immunofluorescence microscopy analysis using rat CSF and brain tissue sections, an increased accumulation of pneumococci was detected over time in the CSF, and because of a loss of drainage between CSF and brain interstitial space, such bacterial accumulation was not observed in the brain parenchyma. Western blot analysis for Iba1, TMEM119 and IFN-Ɣ in rat brain homogenates and NSE in serum showed increased neuroinflammation and neuronal damage in the brain over time during pneumococcal infection. Neurological impairment upon neuronal cell damage caused by meningitis with a malfunctioning glymphatic system was also demonstrated through open-field behavioral tests comparing rats from sham and meningitis groups. Lastly, protein expression analysis of AQP4 revealed no differences in AQP4 between the brains of the rats from the meningitis group and those from the sham non-infected rats. Importantly, confocal microscopy analysis showed a detachment of the astrocytic end feet from the BBB vascular endothelium with consequent misplacement of AQP4-water channels.

**Conclusions:** These findings clearly indicate that pneumococcal meningitis decreases the glymphatic system’s functionality, increasing the neurotoxic waste debris in the brain ultimately leading to brain-wide neuroinflammation and neuronal damage. Finally, our results clearly showed that during pneumococcal meningitis, the glymphatic system does not function because of a detachment of the astrocytic end feet from the BBB vascular endothelium, which leads to a misplacement of AQP4 with consequent the loss of the AQP4-water channel’s functionality.

## Background

*Streptococcus pneumoniae* (the pneumococcus) remains the most significant pathogen responsible for community-acquired bacterial meningitis, a life-threatening inflammation of the meninges surrounding the brain and spinal cord caused by a bacterial infection of the Central Nervous System (CNS) [1]. Importantly, *S. pneumoniae* is the most frequent pathogen associated with bacterial meningitis in both children and adults [2, 3]. In children who survive from an episode of pneumococcal meningitis, the persistent cognitive impairment in low-resource countries is estimated to occur in 4 to 41% [4]. In adults who survived from pneumococcal meningitis in high-resource countries, cognitive impairment is estimated to occur in 32% of survivors [4, 5]. The inflammation triggered by the host immune response leads to the recruitment of peripheral immune cells into the brain and the cerebrospinal fluid (CSF) in a tentative to eliminate the invasive pathogens [6]. The inflammatory mediators which are released in the brain during the infection include cytokines, chemokines, reactive oxygen and nitrogen species, and neurotoxic molecules that contribute to the activation of the microglial cells, initiating neuroinflammation [7, 8, 9]. All these waste products, including host-derived debris, and the invading pathogen itself along with any secreted toxins, are released in the CSF and in the brain parenchyma and must be efficiently cleared from the brain to regain CNS homeostasis. The glymphatic system is responsible for the fluid exchange of the CSF and interstitial fluid, shunting CNS-derived molecules and immune cells from the CNS and meninges to the draining lymph nodes [10]. When the fluid dynamics of the glymphatic system are disrupted, toxic waste accumulates in the brain, further exacerbating the inflammation and interfering with the disease recovery. This study hypothesized that pneumococcal meningitis impaired the glymphatic system’s functionality, decreasing neurotoxic waste clearance in the brain. To test our hypothesis, Wistar rats were subjected to pneumococcal meningitis or artificial-CSF (sham), and the glymphatic system function in the brain and the peripheral circulatory system was analyzed. Intracisternal administration of pneumococci in the CSF progressively caused decreased drainage of CSF into the brain parenchyma, as well as diminished fluid return into the peripheral circulation. We observed a significant accumulation of pneumolysin and pneumococcal capsule localized in the CSF compartment rather than in the brain parenchyma. Neuroinflammation and neuronal damage progressed over time in the brain due to the infection. Lastly, contributing to the malfunctioning of the glymphatic system, we observed during bacterial meningitis astrocytic end feet detachment from the BBB vascular endothelium, which leads to the misplacement of AQP4-water channels that causes the interruption of exchange between the CSF and the brain interstitial space.

## Results

### Impairment of glymphatic system’s functionality during pneumococcal meningitis

Using an experimental meningitis rat model with intracisternal administration of EBA in combination with serotype 3 *S. pneumoniae*, we investigated the loss of functionality of the glymphatic system during CNS pneumococcal infection (Supplementary Figure S1). We measured the EBA levels in the serum of animals subjected to pneumococcal meningitis. At 4, 24, and 72 hours of infection, the meningitis group presented a decrease of the EBA levels in the serum compared with the sham control group demonstrating that the animals presented an impairment of the glymphatic system (Figure 1A). We then evaluated whether the animals presented impairment of the glymphatic system’s functionality by measuring the EBA content retained in the brains of animals subjected to pneumococcal meningitis. As a result, the EBA levels in the meningitis group’s brains were significantly higher than the ones observed in the sham group at all three time-points (Figure 1B).

**Figure 1.**
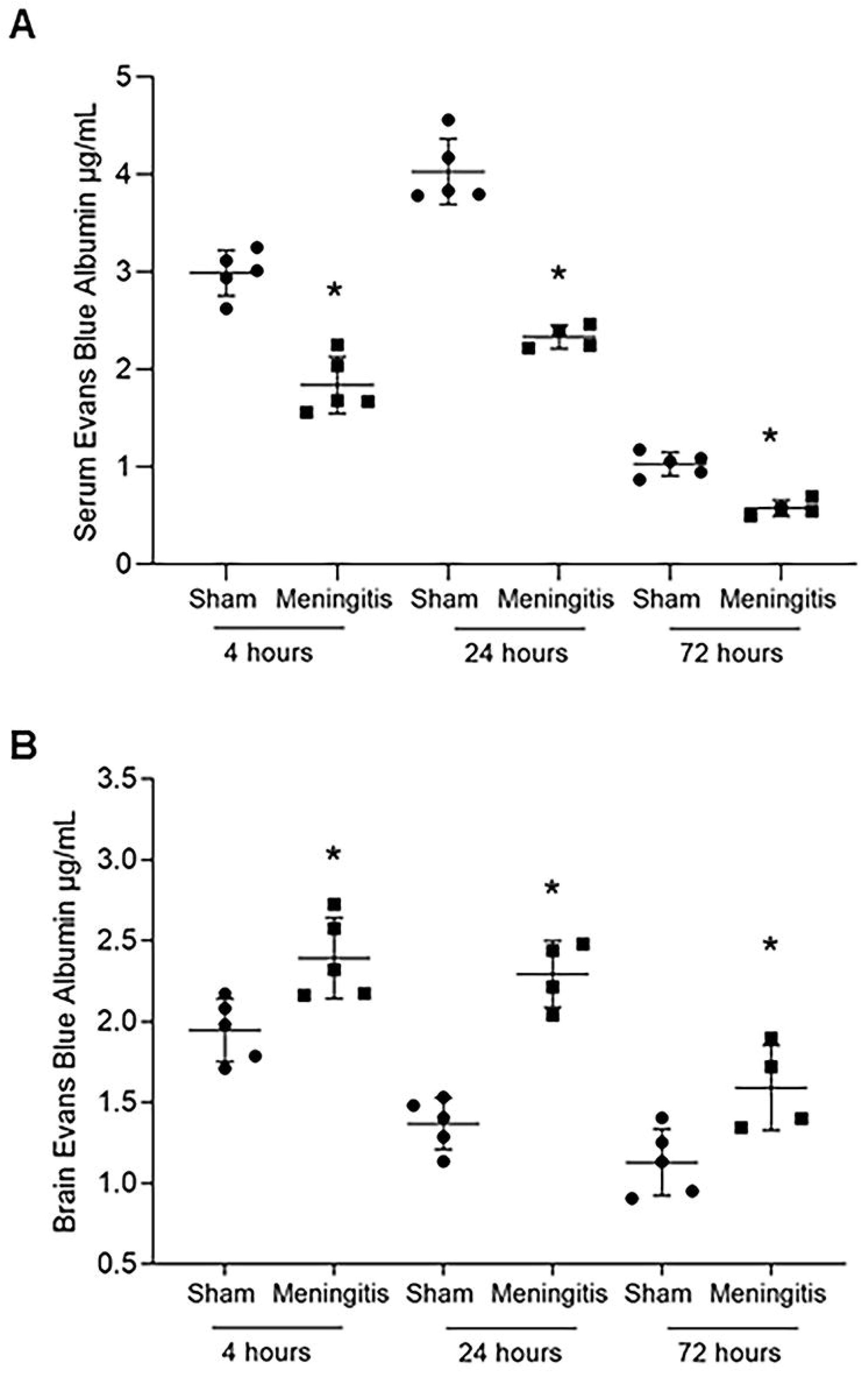
Demonstration of impaired glymphatic system’s functionality during experimental pneumococcal meningitis *in vivo*. Transit of EBA from cisterna magna to serum (A) and from cisterna magna to the brain (**B**) of adult Wistar rats at 4, 24, and 72 hours after induction of pneumococcal meningitis. Data are shown as mean and standard deviation. Statistically significance is shown when compared to the control group (n = 5), * = *p* < 0.05.

### Accumulation of bacterial components in the cerebrospinal fluid but not in the brain parenchyma

To further study the loss of functionality of the glymphatic system during experimental pneumococcal meningitis, we evaluated the presence of pneumococci in the CSF of the rats subjected to pneumococcal meningitis to assess whether an accumulation of bacteria and/or bacterial products in the CSF would occur, due to decreased CSF drainage into the brain parenchyma. Protein levels of pneumolysin (Ply), the main cytotoxin released by pneumococci, were measured by western blotting, and a significant increase of Ply levels was detected in the CSF of the rats from 4 up to 72 hours of pneumococcal infection (Figures 2A-C, and Supplementary Figure S2A). In addition, we also performed immunofluorescence microscopy analysis of the CSF samples, and we observed a significant increase in the fluorescent signal for serotype 3 polysaccharide capsule over time from 4 to 72 hours (Figures 3A and 3B); notably, the non-infected CSF samples did not show any fluorescence signal for pneumococcal capsule (Figures 3A and 3B). Overall, an evident accumulation of bacterial components was detected in the CSF of the rats during pneumococcal infection over time. In line with this accumulation of Ply and capsule, an increase of bacterial viability in the CSF was also detected (Supplementary Table S1). On the other hand, we could not observe the same trend when we analyzed the presence of bacteria in the brain parenchyma. In fact, western blot analysis using brain homogenates revealed that Ply expression levels did not increase over time (Figures 4A-C, and Supplementary Figure S2B), and immunofluorescence microscopy analysis of brain tissue sections showed no accumulation over time of the fluorescent signal of serotype 3 polysaccharide capsule in the brain parenchyma (Figures 5A and 5B). To assess the accumulation of the capsule fluorescent signal over the brain tissue section, we first assessed whether the brain tissue sections from all rats analyzed, both from sham and meningitis groups, had similar tissue surface areas; whole brain tissue sections were imaged using autofluorescence under UV light (Figure 5A), and no significant differences were observed among brain tissue sections from different rats (Supplementary Figure S3). These results can be explained by the loss of functionality of the glymphatic system, which caused retained accumulation of bacterial components in the CSF, and because of the impaired flux from the CSF into the brain tissue, bacterial components did not accumulate in the brain parenchyma as much as in the CSF.

**Figure 2.**
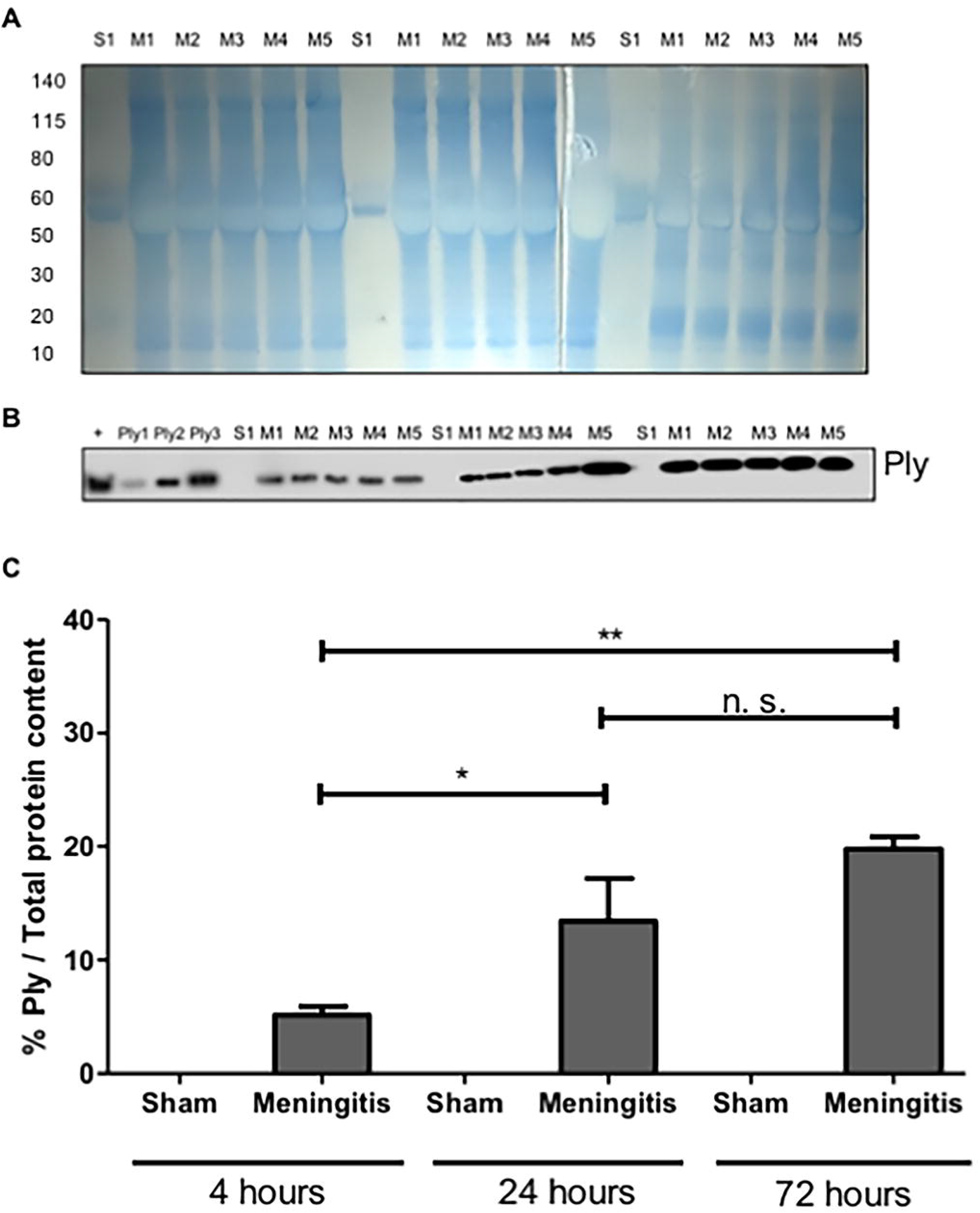
Accumulation of pneumococcal pneumolysin in the cerebrospinal fluid as a consequence of an impaired functionality of the glymphatic system. The total protein content of cerebrospinal fluid (CSF) samples was first measured after InstantBlue Coomassie staining post-SDS-page electrophoresis, S = Sham, M = Meningitis, numbers (1-5) refer to the rat number in the group (Sham or Meningitis) per each time-point (A). Then the same volume of the same samples was used for western blot analysis for the detection of Ply, a lysate of serotype 3 *S. pneumoniae* was used as a positive control (+) together with three serial amounts of purified Ply (Ply1 = 0,0005 µg, Ply2 = 0,005 µg, Ply3 = 0,05 µg in loaded volumes of 10 µl) (**B**). The % of Ply/Total protein content was finally calculated using Image J; data are shown as mean and standard deviation, * = *p* < 0,05, ** = *p* < 0,01, n. s. = non-significant; since sham rats were not infected with pneumococci and therefore did not have any Ply, as a negative control, one CSF sample from sham group for each time point was analyzed, for a total of three sham (non-infected) samples analyzed (**C**).

**Figure 3.**
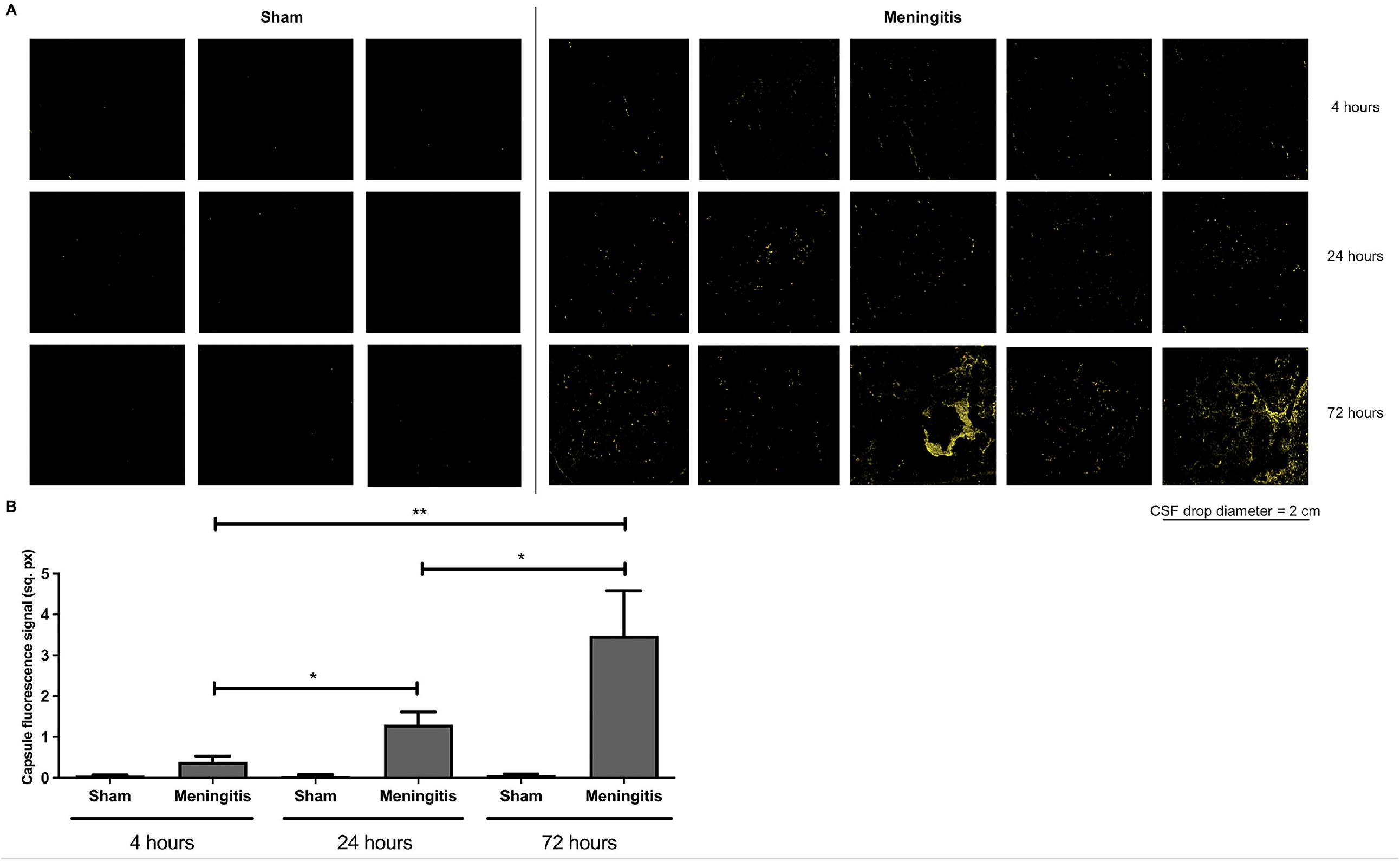
Accumulation of pneumococcal polysaccharide capsule in the CSF due to a malfunctioning glymphatic system during pneumococcal meningitis. 10 µl-CSF drops dried on microscope glass slides stained for serotype 3 polysaccharide capsule (red), the surface area of the fluorescence images is within the yellow border (after the function “Edit selection” of Image J to measure the surface area of the fluorescence signal) to enhance the contract of the fluorescence signal; per each time-point, CSF samples from three rats of the sham group were analyzed in order to have a broad assessment of the unspecific signal from non-infected CSF samples (**A**). Quantification of the fluorescence signal of the pneumococcal capsule stained in Figure 3A, data are shown as mean and standard deviation, * = *p* < 0.05, ** = *p* < 0.01 (B).

**Figure 4.**
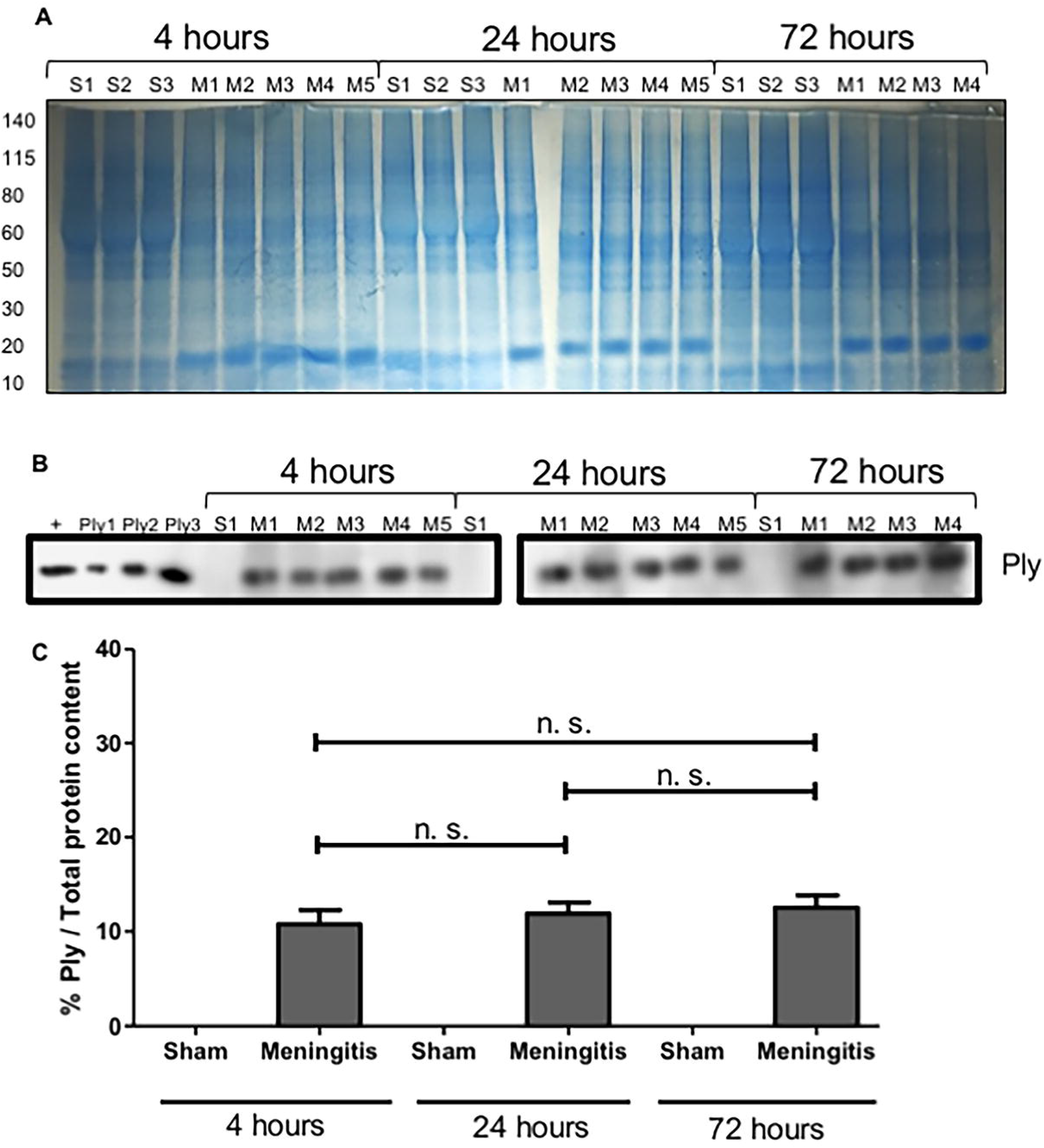
Presence but no accumulation of Ply in the brain homogenates of rats affected by pneumococcal meningitis. The total protein content of brain homogenate samples was first measured after InstantBlue Coomassie staining post-SDS-page electrophoresis, S = Sham, M = Meningitis; numbers (1-5) refer to the rat number in the group (Sham or Meningitis) per each time-point, due to excessive bleeding one brain (M5) from the 72 hour-time-point was not used for this analysis (**A**). Then the same volume of the same samples was used for western blot analysis for the detection of Ply, a lysate of serotype 3 *S. pneumoniae* was used as a positive control (+) together with three serial amounts of purified Ply (Ply1 = 0,0005 µg, Ply2 = 0,005 µg, Ply3 = 0,05 µg in loaded volumes of 10 µl); as a negative control, brain homogenate samples from three rats of the sham (non-infected) group per each time-point were analyzed (**B**). The % of Ply/Total protein content was finally calculated using Image J; data are shown as mean and standard deviation, n. s. = non-significant (**C**).

**Figure 5.**
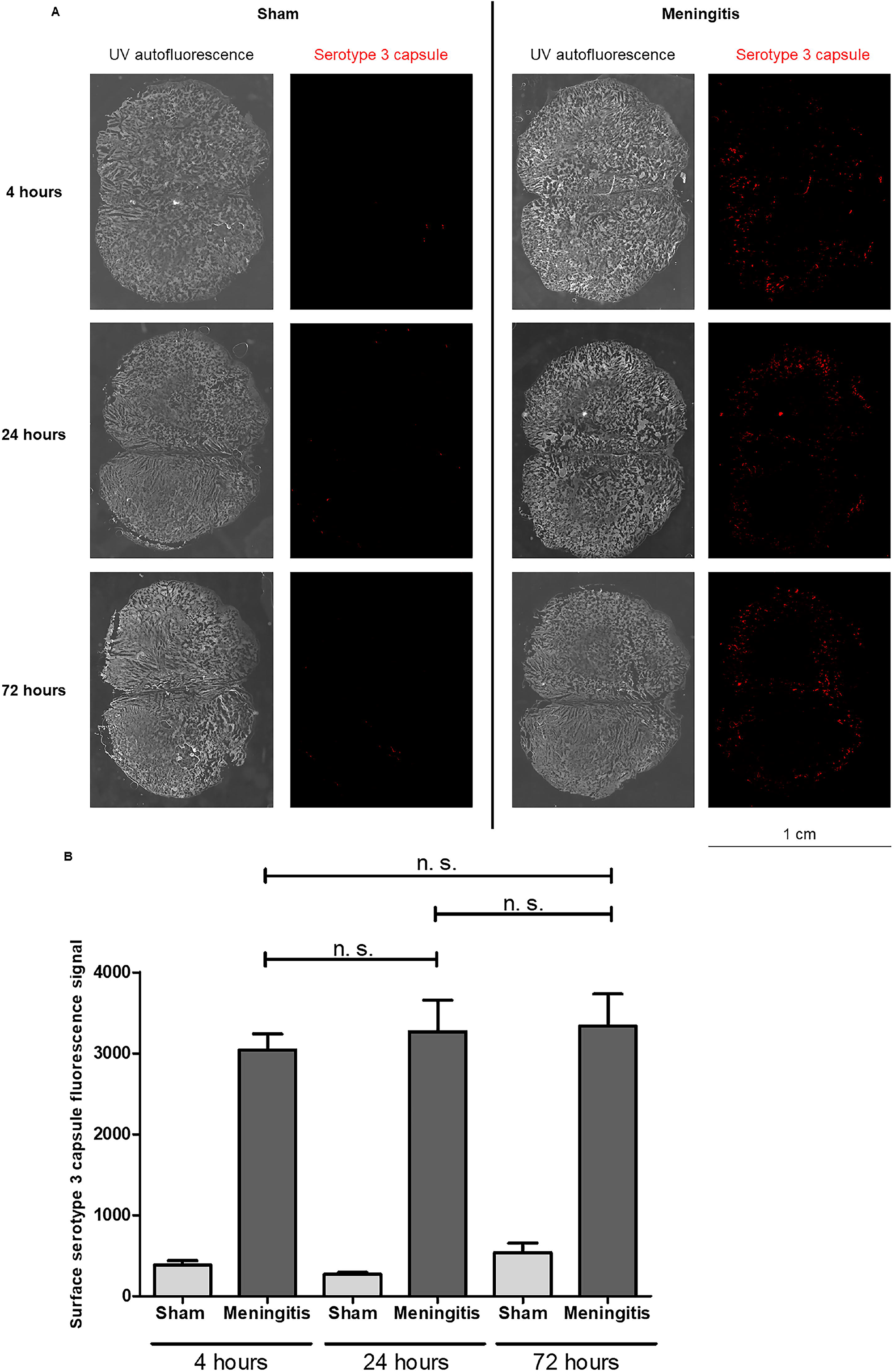
Presence but no accumulation of serotype 3 capsule in the brain parenchyma of rats with pneumococcal meningitis. Brain tissue sections were stained for serotype 3 polysaccharide capsule (red), the whole tissue sections were imaged with autofluorescence under UV light; per each time-point, tissue sections from one rat of the sham group and three rats of the meningitis group and two sections per rat were analyzed in total, one representative image is shown per group (Sham or Meningitis) per each time-point (**A**). Quantification of the fluorescence signal of the pneumococcal capsule stained in Figure 5A, data are shown as mean and standard deviation, n. s. = non-significant (**B**).

### Increased neuroinflammation, neuronal cell damage and impaired neurological functions upon pneumococcal meningitis with a malfunctioning the glymphatic system

As a consequence of an accumulation of bacteria and bacterial components in the CSF, we analyzed the consequent neuroinflammatory status in the brain. We analyzed by western blot the protein levels of ionized calcium-binding adaptor molecule 1 (Iba1), microglia, and macrophage-specific marker [8], using brain homogenates. A significant increase of Iba1 levels was observed in the brains of the rats at 24 and 72 hours compared to the 4 hour-time-point of infection (Supplementary Figures S4A and S4B). This result points towards both an activation of microglia in the brain and an increased infiltration of macrophages into the brain over the course of the intracisternal pneumococcal infection. To further confirm the enhanced inflammatory process during pneumococcal meningitis with an impaired glymphatic system’s functionality, we also measured by western blot analysis the levels of interferon (IFN)-Ɣ, a cytokine with key role of potentiating the pro-inflammatory signaling by priming macrophages and microglia during infections [11, 12]. We observed a prominent increase of IFN-Ɣ in the brain homogenates from 4 up to 72 hours of pneumococcal infection (Figures 6A and 6B). IFN-Ɣ can also be secreted by activated microglia upon neuroinflammation and brain infections, as previously described [13, 14]. Therefore, to assess a local neuroinflammatory response specific for microglia, we have also measured by western blot the expression of the microglial-specific marker TMEM119 in the brain tissue of the rats at different time-points of pneumococcal meningitis. it was recently described that TMEM119 is downregulated in microglia during neuroinflammation and brain disease [15-19]. In line with what reported in literature, we observed a consistent decrease of TMEM119 expression in brain homogenates from 4 up to 72 hours of pneumococcal infection (Figures 6A and 6C). Quantification analysis of the protein band intensities confirmed, respectively, the significant increase of IFN-Ɣ (Figure 6D) and the significant decrease of TMEM119 (Figure 6E) from 4 to up to 72 hours of pneumococcal infection. Moreover, we have also investigated the degree of neuronal damage upon neuroinflammation. Damaged neurons release neuron-specific enolase (NSE), and detection in the blood is often performed in a clinical setting to quantitatively assess the neuronal damage during brain injuries, such as neuroblastoma in newborns, meningitis, and encephalitis [20-23]. We have analyzed the expression levels of NSE in the serum of the rats from the sham and meningitis groups, and a significant increase of NSE expression levels was detected in the serum of the rats from 4 up to 72 hours of pneumococcal infection (Figures 6F-H), indicating that enhanced neuronal damage has occurred in parallel with the increase of neuroinflammation in case of an impaired glymphatic system’s functionality during pneumococcal meningitis. To further confirm this, we performed immunofluorescence microscopy analysis using brain tissue sections and stained for microtubule-associated protein 2 (MAP2) and Tau, respectively expressed mainly on neuronal dendrites and axons [24, 25], in order to semi-quantify the amount of neuronal synaptic connections. We quantified the fluorescent signal of MAP2 and Tau proteins over the whole area of the brain tissue sections, considering that we have previously assessed that the surface areas of the brain tissue sections do not vary significantly between rats from the sham and meningitis groups (Supplementary Figure S3). The brain tissue sections of rats subjected to pneumococcal meningitis showed a significant decrease of MAP2 and Tau fluorescent signals over time, indicating a reduction of neuronal synaptic connections due to neuronal damage (Figures 7A and 7B). To investigate further this neuronal cell damage caused by pneumococcal meningitis with an impaired glymphatic system’s functionality, we performed open-field behavioral tests comparing the sham and meningitis groups. Since behavioral tests require longer times, new animal experiments were conducted in which Wistar rats were intracisternal injected with *S. pneumoniae* and then treated with antibiotic in order for the animals to sustain an infection for ten days. At 10 days the animals were subjected to the open-field task to investigate habituation memory after meningitis induction with a proven impaired glymphatic system. In the sham group, we observed significant differences between training and testing sessions, as measured by the number of crossings and rearings. On the other hand, in the meningitis group we did not observe any difference between training and testing sessions, indication of a consistent impairment of habituation memory (Figure 7C).

**Figure 6.**
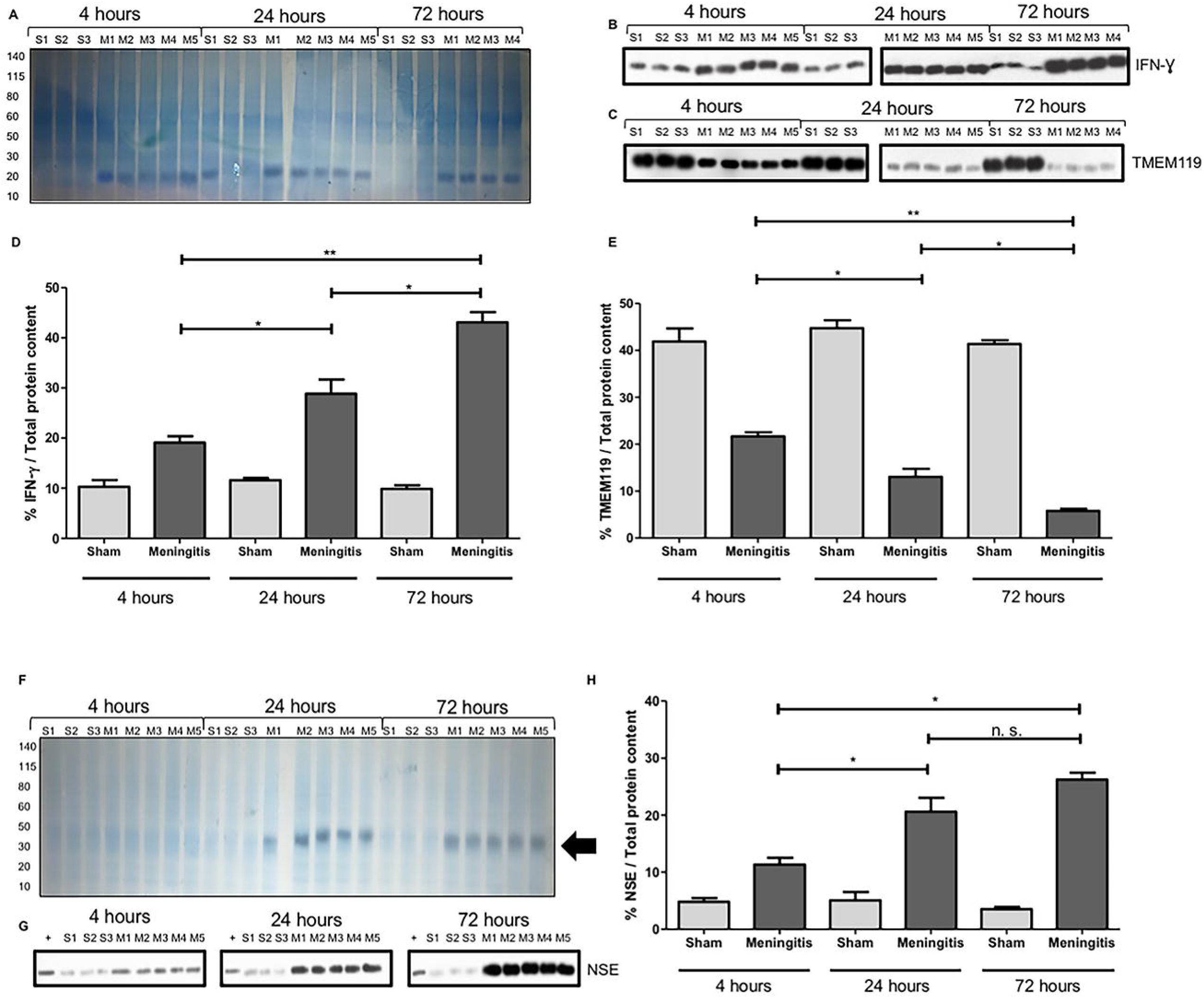
Neuroinflammation and neuronal damage increase over time during pneumococcal infection with a malfunctioning glymphatic system. The total protein content of serum samples was first measured after InstantBlue Coomassie staining post-SDS-page electrophoresis, S= Sham, M= Meningitis; numbers (1-5) refer to the rat number in the group (Sham or Meningitis) per each time-point (**A**). Western blot detection of TMEM119 and IFN-Ɣ was performed using brain homogenate samples of rats from non-infected sham and meningitis groups; since both TMEM119 and IFN-Ɣ are respectively microglial and microglial/macrophage markers present in the brain independently from the infection, brain homogenates from three sham (non-infected) mice were analyzed; due to excessive bleeding one brain (M5) from the 72 hours, time-point was not used for this analysis (**B, C**). The % of TMEM119/Total protein content and IFN-Ɣ/Total protein content were finally calculated using Image J, and data are shown as mean and standard deviation, ** = *p* < 0,01, * = *p* < 0,05; as a negative control, brain homogenate samples from three rats of the sham (non-infected) group per each time-point were analyzed; numbers (1-5) refer to the rat number in the group (Sham or Meningitis) per each time-point (the total protein content was measured after Coomassie staining shown in Figure 6A). (**D, E**). The total protein content of serum samples was first measured after InstantBlue Coomassie staining post-SDS-page electrophoresis, S= Sham, M= Meningitis; numbers (1-5) refer to the rat number in the group (Sham or Meningitis) per each time-point, black arrow points towards an enhanced intensity of the Coomassie staining around the molecular weight of NSE (around 40 kDa) particularly evident for the meningitis group at 24 and 72 hours (**F**). Then the same volume of the same samples was used for western blot analysis for the detection of NSE, a lysate of differentiated neurons from SH-SY5Y cells was used as a positive control (+) (**G**). The % of NSE/Total protein content was finally calculated using Image J, and data are shown as mean and standard deviation, * = *p* < 0,05, n. s. = non-significant (**H**).

**Figure 7.**
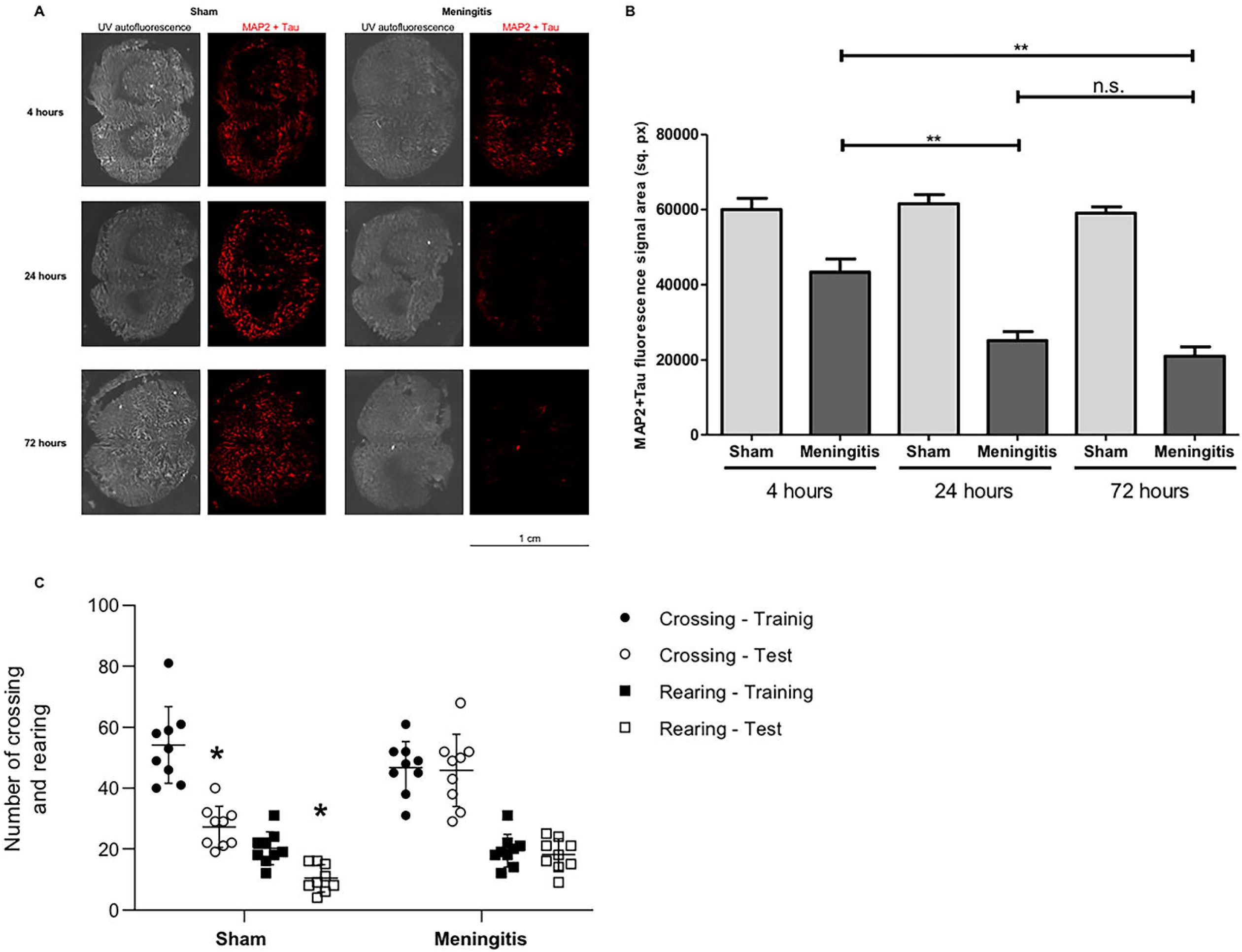
Loss of neuronal synaptic connections and consequent impairment of neurological functions during pneumococcal meningitis with a malfunctioning glymphatic system. Brain tissue sections were stained for a combination of MAP2 and Tau proteins (red), the whole tissue sections were imaged with autofluorescence under UV light; per each time-point, tissue sections from three rats of the sham group and three rats of the meningitis group and two sections per rat were analyzed in total, one representative image is shown per group (Sham or Meningitis) per each time-point (**A**). Quantification of the fluorescence signal of MAP2+Tau stained in Figure 5A, data are shown as mean and standard deviation, ** = *p* < 0.01, n. s.= non-significant (**B**). Open-field task 10 days after pneumococcal meningitis induction; data from the open field task were reported as mean and standard error of mean values, and analyzed by paired Student’s t-tests (n = 9 animals per group). * = p < 0.05 vs. training session.

### Loss of astrocytic interaction with the BBB vascular endothelium and disruption of AQP4-water channels

AQP4 is a water channel present on the end feet of perivascular astrocytes, which are in contact with the vascular endothelium of the BBB, and AQP4-water channels regulate the solute transport of the glymphatic system [26-28]. It was recently reported that AQP4 levels are not altered in the brain cerebral cortex during pneumococcal meningitis [29]. In line with these recent findings, our western blot analysis using brain homogenates clearly showed that upon experimental pneumococcal meningitis, AQP4 protein levels are not altered comparing brain tissue of the rats from sham and meningitis groups over time (Figures 8A and 8B). We then performed high-resolution immunofluorescence microscopy analysis using brain tissue sections stained for the vascular endothelium of the BBB using Lycopersicon Esculentum tomato lectin DyLight 594 [30], and astrocytes using anti-glial fibrillary acidic protein (GFAP) antibody. Clear signs of astrogliosis were observed over time during pneumococcal meningitis infection, in particular activated astrocytes, which upon neuroinflammation retracted their cellular processes, showed a progressive detachment of the astrocytic end feet from the BBB vascular endothelium, likely leading to the loss of function of AQP4-water channels (Supplementary Figure S5). The increased detachment of the astrocytic end feet from the brain vascular endothelium was further confirmed by the co-localization analysis performed by Image J, which showed the connection between astrocytic end feet and the BBB endothelium is lost over time (Figure 8C). To assess the misplacement of AQP4-water channels as a consequence of the astrocytic cellular process retraction, we have also performed confocal microscopy analysis combined with 3D reconstruction modeling. It is evident that during the course of pneumococcal infection, the progressive retraction of the astrocytic cellular processes causes the displacement of the AQP4 which loses its natural localization connecting the astrocytic end feet with the BBB vascular endothelium (Figure 8D). Furthermore, while in the brain of sham control animals with a functioning glymphatic system the AQP4 expression is homogeneously expressed along the area between astrocytic end feet and BBB vascular endothelium, in the brain of meningitis-affected rats in which the glymphatic system is malfunctioning, the AQP4 expression is focal on astrocytic cells far without any co-localization with the BBB endothelium (Figure 8D).

**Figure 8.**
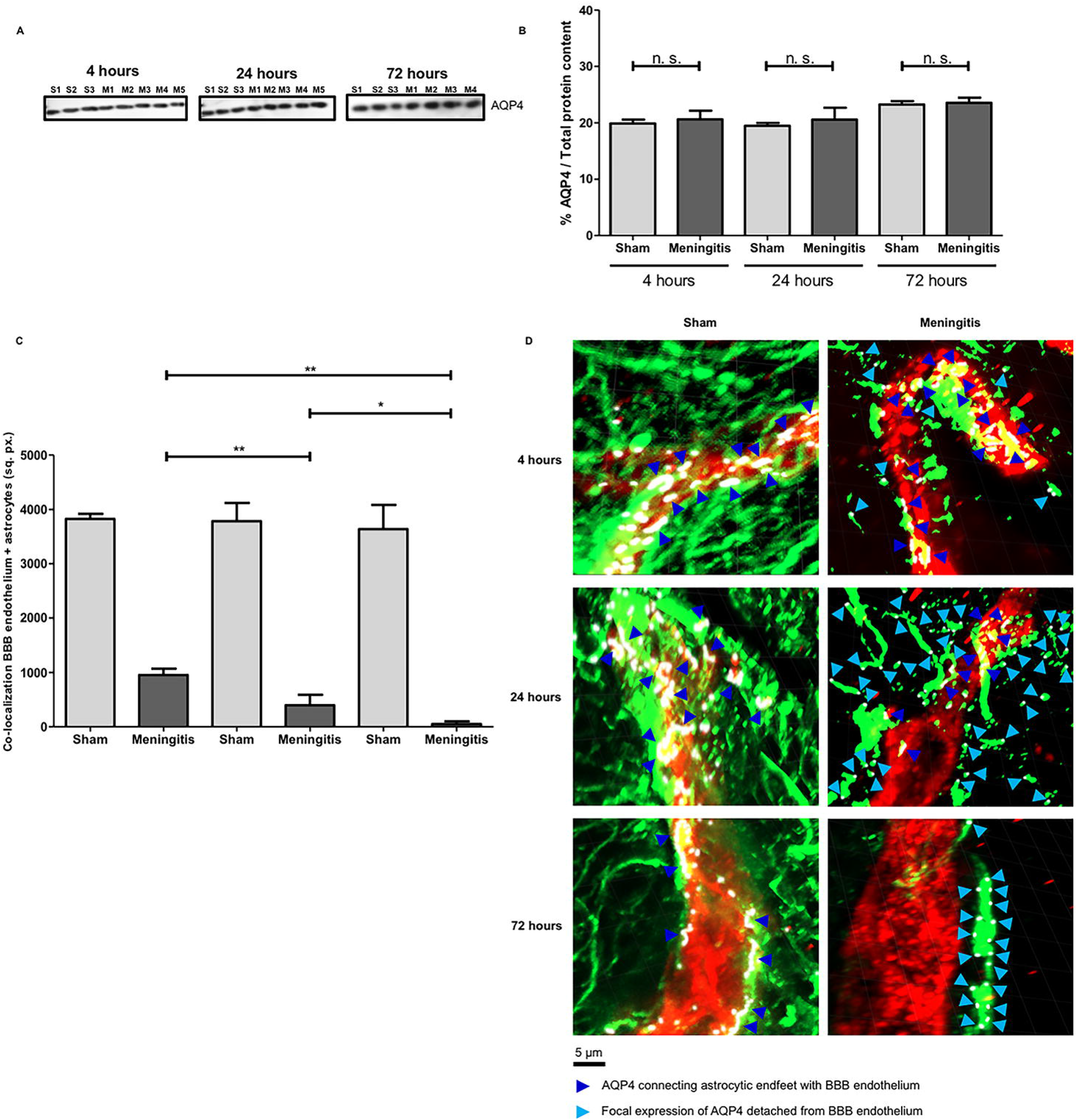
Loss of glymphatic system’s functionality is not due to changes in AQP4 expression levels but to a detachment of astrocytic end feet from BBB vascular endothelium. Western blot detection of AQP4 in brain homogenate samples from sham and meningitis groups; since AQP4 is expressed by astrocytes independently from the infection, brain homogenates from three sham (non-infected) mice were analyzed; due to excessive bleeding one brain (M5) from the 72 hour-time-point was not used for this analysis (**A**). The % of AQP4/Total protein content was finally calculated using Image J, and data are shown as mean and standard deviation, n. s. = non-significant (The total protein content was measured after Coomassie staining shown in Figure 4A) (**B**). High-resolution immunofluorescence microscopy analysis of BBB vascular endothelium in the cerebral cortex stained with Lycopersicon Esculentum tomato lectin DyLight 594 (red) and astrocytes stained with mouse anti-GFAP antibody combined with Alexa Fluor goat anti-mouse 488 (green). Brain tissue sections from sham rats clearly show astrocytes in close proximity around the brain vasculature in support of the BBB; in contrast, brain tissue from meningitis-affected rats showed a progressive detachment overtime of the astrocytic end feet from the vascular endothelium; the white double-arrows point towards the detaching astrocytic end feet. Brain tissue sections from three sham mice and five (four for the 72 hour-time-point) were analyzed per time point, four brain tissue sections per rat were used for the microscopy analysis (**C**). Quantification of the area of co-localization between the BBB vascular endothelium (in red, shown in Supplementary Figure S5) and the astrocytes (in green, shown in Supplementary Figure S5) measured with Image J, data are shown as mean and standard deviation, * = *p* < 0.05, ** = *p* < 0.01. (**D**) Confocal microscopy analysis of BBB vascular endothelium in the cerebral cortex stained with Lycopersicon Esculentum tomato lectin DyLight 594 (red), astrocytes stained with mouse anti-GFAP antibody combined with Alexa Fluor goat anti-mouse 488 (green), and AQP4 stained with rabbit anti-AQP4 antibody combined with Alexa Fluor goat-anti rabbit 647 (purple, assigned a white color using the imaging program ZEN lite); dark blue arrows point towards AQP4 fluorescent signal connecting astrocytic end feet with BBB vascular endothelium is, while light blue arrows point towards focal expression of AQP4 on astrocytes detached from the BBB endothelium; six brain tissue sections from three sham rats and five (four for the 72 hour-time-point) from the meningitis group were analyzed per time point, three images per section were taken, the displayed images are representative of each group; plane images were angled of 30 degrees from horizontal position on the Z axis using Imaris.

## Discussion

The BBB, in the CNS, allows passage of O2, CO2, and glucose as a demand for brain cell metabolism but prevents unregulated protein and fluid entry into the brain [31]. However, the interstitial space of the brain still receives soluble waste products as a consequence of its elevated metabolism [32]. The glymphatic system maintains the balance by supporting the delivery of fluid and interstitial solutes, including immune cells and macromolecules from the brain to the CSF egress routes to draining in the lymph nodes [33]. The glymphatic dysfunction is characterized by a reduced exchange between CSF and interstitial fluid (ISF), leading to an accumulation of waste products. Many diseases have been associated with glymphatic system dysfunction, such as Alzheimer’s disease, Parkinson’s disease, and other neurodegenerative diseases [34-36]. The BBB transport and glymphatic clearances are interdependent mechanisms, and the dysfunction of these mechanisms can difficult the solute clearance [36]. The BBB permeability or disruption facilitates the migration of peripheral immune cells and inflammatory mediators into the CSF, increasing the production of cytokines, chemokines, reactive oxygen and nitrogen species, and neurotoxic molecules that contribute to glial cell activation and consequent neuroinflammation [6, 7, 8, 9]. Pre-clinical studies of bacterial meningitis have demonstrated that the BBB breakdown is associated with cognitive impairment [37, 38], and when the proteolytic enzymes like matrix metalloproteinases (MMPs) were blocked cognitive dysfunction could be prevented [39, 40]. This study evaluates the glymphatic function in an animal model of pneumococcal meningitis. The meningitis group presented an accumulation of the EBA in the brain compared with the control group, demonstrating that the glymphatic function was affected by meningitis. The EBA has not drained adequately from the CSF to the lymphatic nodes accumulating it in the brain of the animals. To confirm this result, we also evaluate the EBA levels in the bloodstream of the animals. The control group presented an increase in EBA levels in the serum compared with the meningitis group, demonstrating that the meningitis group was unable to drain the EBA from the CSF to the bloodstream. As a consequence of the loss of solute drainage between the CSF and the brain parenchyma, most of pneumococcal components remained accumulated in the CSF.

Neuronal injury is frequently observed upon neuroinflammation. Damage of neurons can be caused both by direct interaction of the bacteria or bacterial components, like pneumolysin, with neurons [41], and by the damaging effect of pro-inflammatory compounds released during the neuroinflammation process [42]. In line with this, associated with a loss of glymphatic system functionality during pneumococcal meningitis pathogenesis, we showed that neuroinflammation and neuronal injury increase overtime during the infection. Therefore, our data strongly suggest that the glymphatic system dysfunction could be one more factor associated with long-term cognitive impairment in meningitis survivors. The bacterial meningitis survivors’ patients present an increased risk of triggering neurocognitive impairment. In a children’s birth cohort, meningitis in early-life was associated with neurocognitive, educational, and psychological difficulties during childhood and early adolescence [43]. Another study also demonstrated that meningitis during childhood increases the risk of schizophrenia in adulthood [44]. In adults, neurologic sequelae of pneumococcal meningitis in high-resource countries presented a rate of 32% of cognitive impairment, 4% of hydrocephalus, 31% seizures, 22 to 69% hearing loss, and 11 to 36% of focal deficits [4]. The connections between BBB disruption, glymphatic dysfunction, and cognitive impairment in meningitis survivors could be a new avenue to investigate mechanisms to prevent these events and re-establish an everyday life for these patients committed to bacterial meningitis.

AQP4-water channels are a fundamental component of the glymphatic system, facilitating the solute transport from the perivascular subarachnoid space and the brain parenchyma [26-28]. Upregulation of AQP4 expression levels was previously described upon neuroinflammation in neurodegenerative diseases, such as Alzheimer’s disease [45] and multiple sclerosis [46]. On the other hand, Pavan and collaborators have recently observed that upon neuroinflammation caused by a pneumococcal infection in the CNS, AQP4 expression levels are not significantly altered compared to a non-infected situation. Our results go in line with Pavan et al [29], as we did not observe increased AQP4 expression levels in the brains from the meningitis-affected rats compared to the brains from the sham control mice. Significantly, what affects the solute transport of the glymphatic system during pneumococcal meningitis is not the altered AQP4 expression, but rather the disruption of the AQP4-water channels due to a detachment of the astrocytic end feet from the BBB vascular endothelium. This cellular process is a straightforward consequence of the morphological changes of astrocytes that, during neuroinflammation, undergo astrogliosis, in which the cell body becomes rounder and more prominent, and the cellular processes become ticker and shorter [47, 48]. The shortening of the astrocytic cellular processes causes the detachment of the astrocytic end feet from the BBB vascular endothelium; such detachment causes the misplacement, and consequent loss of function, of the AQP4-water channels which usually is present across the interspace between the brain vascular endothelium and astrocytic end feet.

## Conclusions

In conclusion, in this study we showed that during experimental pneumococcal meningitis, a loss of functionality of the glymphatic system occurs with a consequent accumulation of pneumococcal components, such as the cytotoxin Ply and the bacterial polysaccharide capsule used as in this study as markers for the presence of *S. pneumoniae*, in the perivascular subarachnoid space occupied by CSF. Such accumulation of bacteria leads to a dramatic increase in neuroinflammation and neuronal injury; therefore, the loss of glymphatic system’s functionality can play a crucial role in determining the neurological sequelae which often occur in patients who survive from bacterial meningitis [4, 41]. Astrocytes are a fundamental component of support for the BBB, with astrocytic end feet in close contact with the BBB vascular endothelium [49]. Here, we reported for the first time that the impairment of solute transport between perivascular subarachnoid space and the brain parenchyma occurs when a detachment of astrocytic end feet from the BBB vascular endothelium occurs, with consequent misplacement and likely loss of the physiological function of AQP4-water channels of solute transport within the glymphatic system.

## Material and Methods

### Infecting organisms to induce meningitis

The serotype 3 *S. pneumoniae* was cultured in 5 mL of Todd Hewitt Broth®, diluted in fresh medium, and then grown to the logarithmic phase. The culture was centrifuged for 10 minutes at 1200 rpm and resuspended in sterile saline to a concentration of 5 × 10^9^ colony-forming units (CFU) [40].

### Animal model of meningitis

Male Wistar rats (8-weeks-old, 200 to 250 g body weight), obtained from our breeding colony and used for the experiments, where remained housed in a cool, on a 12-hour light/dark cycle, at a temperature of 23°C ± 1°C, with food and water always available *ad libitum*. Scoring of the clinical symptoms of the rats during pneumococcal infection was performed according to the research protocol was approved by the Animal Care and Experimentation Committee of UNESC 93/2019, Brazil, and realized following the National Institutes of Health Guide for the Care and Use of Laboratory Animals (NIH Publication No. 80-23, revised in 1996) (Supplementary Table S1). The animals were anesthetized with an intraperitoneal (i.p.) administration of ketamine (6.6 mg/kg) and xylazine (0.3 mg/kg) [50, 51]. The animals were infected intracisternal with 10 μl of the inoculum containing 5 × 10^9^ CFU/ml of living *S. pneumoniae*. Moreover, the control sham animals were injected with 10 μl of artificial cerebrospinal fluid (aCSF). Meningitis (n = 5) and the control group (n = 5) received fluid replacement after bacterial induction and were then returned to their cages. Meningitis was confirmed by incubating a quantitative culture of 5 μL of CSF at 37°C with 5% CO2 on sheep blood agar [52]. For the 10 day-infection, after 18 hours of meningitis induction, the animals submitted to behavioral test received ceftriaxone (100 mg/kg, i.p., for seven days) twice a day. Ten days after pneumococcal meningitis induction, nine animals from the sham group and nine animals from the meningitis group were submitted to the behavioral test of habituation to the open field.

### Intracisternal injection of Evans blue albumin

Previous reports demonstrated that radiolabeled albumin, and Evans’ blue albumin (EBA), when are injected into the brain parenchyma, migrate and concentrate in perivascular spaces, lymph vessels in the neck, and the cervical lymph nodes [53, 54]. With fluorescent and colorimetric properties, Evans blue binds with affinity to albumin and has been used to assess the permeability of the BBB [55-57]. The 1% solution of Evans blue albumin (EBA) was prepared to add 100 mg of Evans blue (Sigma-Aldrich, St. Louis, MO) and 100 mg of rat albumin (Sigma-Aldrich, St. Louis, MO) diluted in 10 mL artificial cerebrospinal fluid (aCSF) [49]. A 30G dental needle was inserted into one end of a ca 30 cm PE10 tubing and was connected to the other end of the tubing to a 100 μL Hamilton syringe. Then the cannula was filled with 30 μL of EBA solution. In the sequence, the syringe was attached to the micro-infusion pump. The previous anesthetized animal that received the bacterial suspension or aCSF was placed in the stereotaxic on a heating pad, and the head was fixed on the apparatus with the rat’s nose slightly pointed downwards. The cannula needle was inserted (approximately 1 to 2 mm) into the center of the cisterna magna, and it started the injection of EBA using the microinjection syringe pump at a rate of 1 μL per minute for 25 min, resulting in a total volume injected of 25 μL of 1% EBA solution [58].

### Quantification of Evans blue albumin in serum

At 4, 24, and 72 hours after the meningitis induction and EBA solution injection in the cisterna magna, blood was collected in the femoral artery and centrifuged for 10 min at 2000 × g to isolate the serum. The rats were euthanized by decapitation under anesthesia, and the brains were placed in 3 mL of formamide (Sigma-Aldrich®, St. Louis, MO) to allow EBA to diffuse from brain tissue and hold there for 72 h. The brains were removed, and the formamide was stored at -20°C until the spectrophotometry evaluation. The EBA concentration was measured in the serum and the formamide solution. Both samples were evaluated using spectrophotometry (620 nm) [59].

### Behavioral open-field tests

Ten days after pneumococcal meningitis induction, nine animals from the sham group and nine animals from the meningitis group were submitted to the behavioral test of habituation to the open field. After the behavioral study, the rats were euthanized, and the brain was dissected for neurochemical evaluations. The open-field task evaluates motor performance in the training section and non-associative memory in the retention test session. Habituation to an open field was carried out in a 40 × 60 cm open field surrounded by 50 cm high walls made of brown plywood with a frontal glass wall. The floor of the open field was divided into 9 equal rectangles by black lines. Each animal was gently placed on the left rear quadrant and left to explore the arena for 5 min (training session). Following this, the animals were taken back to their home cage and 24 hours later submitted again to a similar open-field session (test session). The crossings number (number of times that the animal crossed the black lines, an assessment of locomotor activity) and rearing movements (exploratory behavior observed in rats subjected to a new environment) were performed in both sessions and were counted. The decrease in the number of crossings and rearings between the two sessions was taken as a measure of the retention of habituation memory [60]. The test was conducted by a person who was blinded to the groups.

### Protein lysate preparation for SDS-page electrophoresis

Brains were homogenized in RIPA buffer containing protease and phosphatase inhibitors using a cell strainer with a 100 µm filter (Falcon). Brain homogenates, as well as CSF, serum samples, and purified Ply, the liquid culture of type 3 *S. pneumoniae* OD600 0.5 were mixed with LDS sample buffer (stock 4X concentrated down to 1X concentration in each sample) (Thermo Fisher Scientific) and boiled at 95°C for 5 minutes.

### Coomassie instant blue staining, western blot analysis, and protein expression quantification

Protein samples were loaded into NuPage Novex 4-12% Bis-Tris SDS-PAGE gels (Thermo Fisher Scientific), electrophoresis was performed using Mini Gel Tank (Thermo Fisher Scientific), and electroblotting was performed on PVDF membranes (Novex, Life Technologies) using Mini Blot Module (Thermo Fisher Scientific). Ply, NSE, Iba1, IFN-Ɣ, TMEM119 and AQP4 were detected using respectively a mouse anti-Ply antibody (Abcam), a rabbit anti-NSE antibody (Merck), a goat anti-Iba1antibody (Abcam), a rabbit anti-IFN-Ɣ antibody (Abcam), a rabbit anti-TMEM119 antibody (Thermo Fisher) and a rabbit anti-AQP4 antibody (Abcam); all primary antibodies were diluted 1:1000 in PBS 0.1% Tween (PBS-T); the secondary antibodies used were HRP-conjugated goat anti-mouse, goat anti-rabbit and donkey anti-goat (Thermo Fisher Scientific), all diluted 1:5000 in PBS-T. Protein bands were detected by incubating membranes with ECL Prime Western Blotting Detection Reagents (Cytiva, Thermo Fisher Scientific) and imaged using ImageQuant LAS 4000 (GE Healthcare). For Coomassie stainings, after electrophoresis gels were incubated with InstantBlue Coomassie Protein Stain (Abcam) for 15 minutes and then photographed. The intensity of protein bands on PVDF membranes was measured by Image J as previously described [61]. Percentages of protein expression were calculated as a ratio of the protein band intensity divided by the total protein content (after Coomassie staining) measured by Image J in each specific lane of the gel/membrane, the same volume of protein lysate was loaded onto gels for Coomassie staining to perform the quantification of the total protein content in each sample, and on a separate gel for protein detection by western blot.

### Immunofluorescence stainings

Coronal sections of 20 µm thickness of snap frozen rat brains embedded in Cryomatrix (Thermo Fisher Scientific) were cut and collected on Superfrost Plus slides (Thermo Fisher Scientific). Sections were fixed with acetone for 10 minutes, dried, edges of the sections marked with PAP pen (Avantor, VWR), and incubated with one of the following combinations: 1. Rabbit polyclonal anti-capsule serotype 3 antiserum (SSI Diagnostica) followed by Alexa Fluor 594 goat anti-rabbit (Thermo Fisher Scientific), 2. Mouse anti-MAP2 antibody combined with chicken anti-Tau antibody (Abcam) followed by Alexa Fluor 594 goat anti-mouse (Thermo Fisher Scientific) and Alexa Fluor 594 goat anti-chicken (Thermo Fisher Scientific), 3. Mouse anti-GFAP antibody (Santa Cruz Biotechnology) followed by Alexa Fluor 488 goat anti-mouse (Thermo Fisher Scientific), combined with Rabbit anti-AQP4 antibody followed by Alexa Dluor 647 goat anti-rabbit (Thermo Fisher Scientific) mixed with Lycopersicon Esculentum tomato lectin DyLight 594 (Vector Laboratories). Incubations with primary antibodies were performed for two hours at room temperature, incubations with secondary antibodies were performed for two hours at room temperature in the dark.

For staining of CSF samples, drops of 10 µl were pipetted onto microscope glass slides and let dry at room temperature. Once the CSF drops were completely dry, the edges of each drop were marked with PAP pen and incubated with Rabbit polyclonal anti-capsule serotype 3 antiserum (SSI Diagnostica) for 1 hour at room temperature, followed by Alexa Fluor 594 goat anti-rabbit (Thermo Fisher Scientific) for 1 hour at room temperature in the dark.

All samples were blocked with PBS with 5% bovine serum albumin (BSA) (Sigma Aldrich) for 15 minutes at room temperature before starting the staining procedure. All antibodies were diluted according to the recommendations from the manufacturers using PBS with 1% BSA, and slides were washed in PBS three times for 5 minutes between primary and secondary antibody incubations. Slides were finally mounted using ProLong Diamond Antifade Mountant (Invitrogen, Thermo Fisher Scientific), and coverslips were added.

### High-resolution fluorescence, confocal microscopy imaging and 3D modeling

All stained samples were imaged using a Zeiss Observer.Z1 fluorescence microscope with Orca-Flash4.OLT camera. To acquire the images of the entire CSF 10 µl drops of the brain tissue sections, a 5X objective combined with the function “Tiles” of the imaging program Zen 2 (Zeiss) was used. For imaging with higher magnification of BBB vascular endothelium (tomato lectin) and astrocytes (anti-GFAP antibody followed by Alexa Fluor 488 goat anti-mouse), a 63X objective combined with the function “Stacks” of the imaging program Zen 2 were used, each image taken was a merge of 15-20 stacks imaged. Confocal microscopy analysis was performed using a Zeiss LSM980-Airy2 confocal system; to acquire images, the program ZEN lite (Zeiss) was used, while 3D modeling after image acquisition was performed using Imaris (Oxford Instruments). During the confocal microscopy imaging, each z-stack-image included 15 stacks of a total thickness of 15 µm.

### Quantification of the fluorescence signal and co-localization analysis

Fluorescence signal post-stainings was quantified using the software Image J as previously described [62]. Briefly, all fluorescence images were converted into grayscale images, and then using the function “Threshold", the surface covered by the fluorescence signal was defined with the function “Create Selection” and finally quantified using the function “Measure.” For colocalization analysis, fluorescence signals from tomato lectin and astrocytes (stained with anti-GFAP antibody) were analyzed for co-localization using the “Co-localization” plugin, which turns all the co-localized signals into white color; the surface covered by the co-localized white color was then measured using the function “Measure.”

### Statistical analysis

For statistical analysis related to the quantification of Evans blue albumin in Wistar rats, the software SPSS was used; for the statistical analysis related to all other *ex vivo* analyses, the software Graph Pad (Prism 5) was used. For all multiple comparisons, nonparametric ANOVA was used to assess the presence of differences between the groups, and then a Dunn’s test was applied for pairwise comparisons. For 2-group comparisons, the nonparametric 2-tailed Wilcoxon’s rank-sum test (also known as the Mann-Whitney *U* test) was used. *P* values are indicated in each figure legend.

## Abbreviations

CNS: Central Nervous System
BBB: blood-brain barrier
CSF: cerebrospinal fluid
AQP4: Aquaporin 4
Evans Blue Albumin: EBA
Ply: Pneumolysin
Iba: ionized calcium-binding adaptor molecule 1
IFN: interferon
TMEM: transmembrane protein
NSE=: neuron-specific enolase
MAP2: microtubule-associated protein 2
GFAP: glial fibrillary acidic protein
ISF: interstitial fluid
MMPs: matrix metalloproteinases
CFU: colony forming unit

## Acknowledgments

This work was supported by the Faillace Department of Psychiatry and Behavioral Sciences, McGovern Medical School, The University of Texas Health Science Center at Houston (UTHealth), USA (TB); the Graduate Program in Health Sciences, University of Southern Santa Catarina (UNESC) (TB, JSG, AC, and CJF), Brazil, and the Alzheimer’s Association Grant number AARGDNTF-19-619645 (TB). We thank Dr. Fariba Foroogh from the Department of Medicine Solna at Karolinska Institutet for the help with the cryostat brain tissue cutting, Prof. Birgitta Henriques-Normark from the Department of Microbiology, Tumor and Cell Biology at Karolinska Institutet for the scientific discussions, Dr. Miguel Tofiño Vian from the Department of Neuroscience at Karolinska Institutet for providing a lysate of differentiated neurons from SH-SY5Y cells, the Protein Production Platform at Nanyang Technological University in Singapore for providing the purified Ply, and the Biomedicum Imaging Core Facility for the help during the confocal microscopy analysis and 3D modeling.

## Ethics Declaration

### Animal experiments

Experiments with Wistar rats were approved by the Animal Care and Experimentation Committee of UNESC 93/2019, Brazil, and realized following the National Institutes of Health Guide for the Care and Use of Laboratory Animals (NIH Publication No. 80-23, revised in 1996).

### Consent of publication

All authors have agreed with the publication of the manuscript in case of acceptance after peer-review according to the Open Access policy of the journal.

### Availability of data and material

The datasets used and/or analysed during the current study are available from the corresponding author on reasonable request.

### Author Contributions

F.I., T.B., J.S.G., and S.T. designed the study, wrote the protocols, undertook the experimental analysis, and wrote the first draft of the manuscript. J.S.G., S.S., A.C., R.R.E.S. and D.D. performed the experiments. All authors contributed to and have approved the final manuscript.

