## Supplementary Material for "Dysfunctional glymphatic system with disrupted aquaporin-4 expression pattern on astrocytes causes bacterial product accumulation in the CSF during pneumococcal meningitis"

### Supplementary Information

**Supplementary Figure S1. Representation of EBA injection in the cisterna magna**. The needle was inserted into the center of the cisterna magna, and it started the injection of EBA using the micro-injection syringe pump. The solute is drained from the cerebral parenchyma by the perivenous fluid to the peripheral lymph nodes and circulatory system.

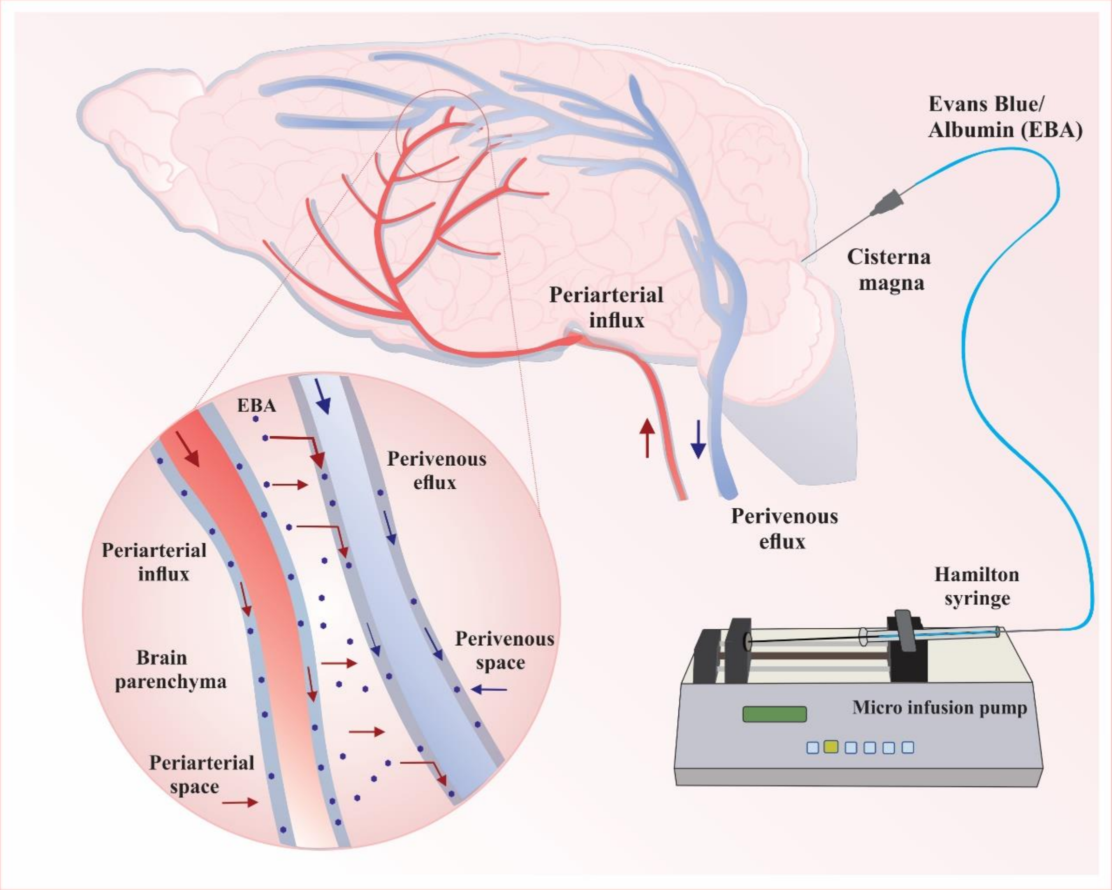

**Supplementary Figure S2. Putative concentrations of Ply in the CSF and brain parenchyma of rats affected by pneumococcal meningitis.** Ply concentrations were calculated by measuring the band intensity of purified Ply at three different serial concentrations (Ply1 = 0,0005 µg, Ply2 = 0,005 µg, Ply3 = 0,05 µg in loaded volumes of 10 µl) using Image J, accordingly the putative concentrations of Ply in the CSF (**A**) and brain homogenate samples (**B**) were calculated.

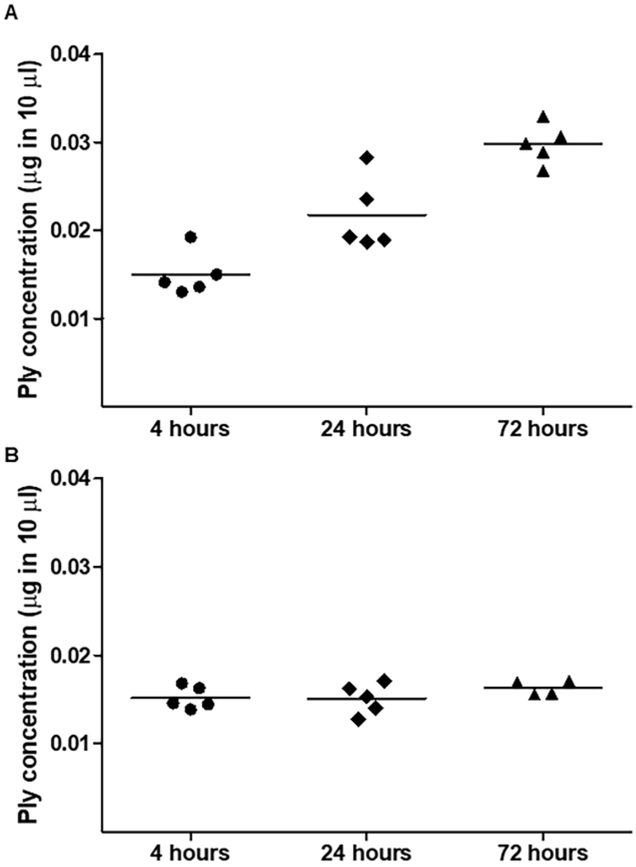

**Supplementary Figure S3. The surface area of the brain tissue sections.** Quantification of the surface area (sq. px. = square pixels) occupied by the brain tissue sections used for immunofluorescence microscopy (results shown in Figure 5); for each rat, two brain sections were analyzed.

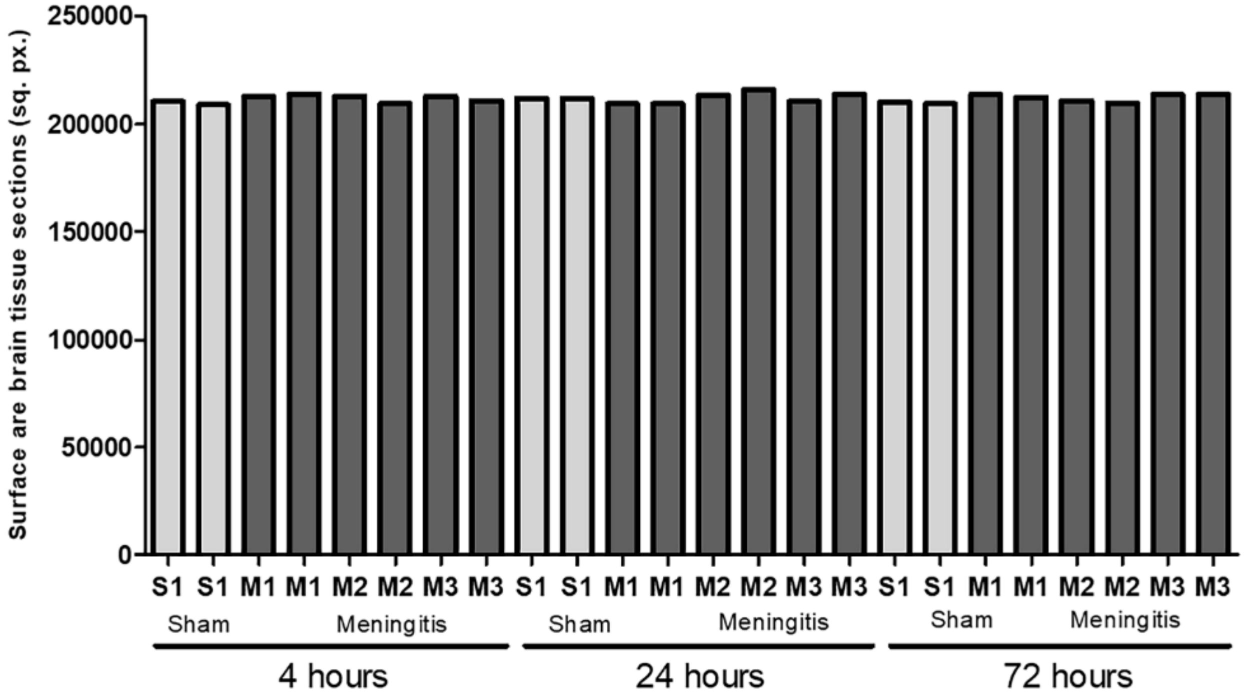

**Supplementary Figure S4. Increase of the macrophage and microglia marker Iba1 during pneumococcal infection with a malfunctioning glymphatic system.** Western blot detection of Iba1 in brain homogenate samples of rats from non-infected sham and meningitis groups; since Iba1 is a microglial/macrophage marker present in the brain independently from the infection, brain homogenates from three sham (non-infected) mice were analyzed; due to excessive bleeding one brain (M5) from the 72 hours, time-point was not used for this analysis (**A**). The % of Iba1/Total protein content was finally calculated using Image J, and data are shown as mean and standard deviation, ** = $p < 0{,}01$, n. s. = non-significant; as a negative control, brain homogenate samples from three rats of the sham (non-infected) group per each time-point were analyzed; numbers (1-5) refer to the rat number in the group (Sham or Meningitis) per each time-point (the total protein content was measured after Coomassie staining shown in Figure 4A)

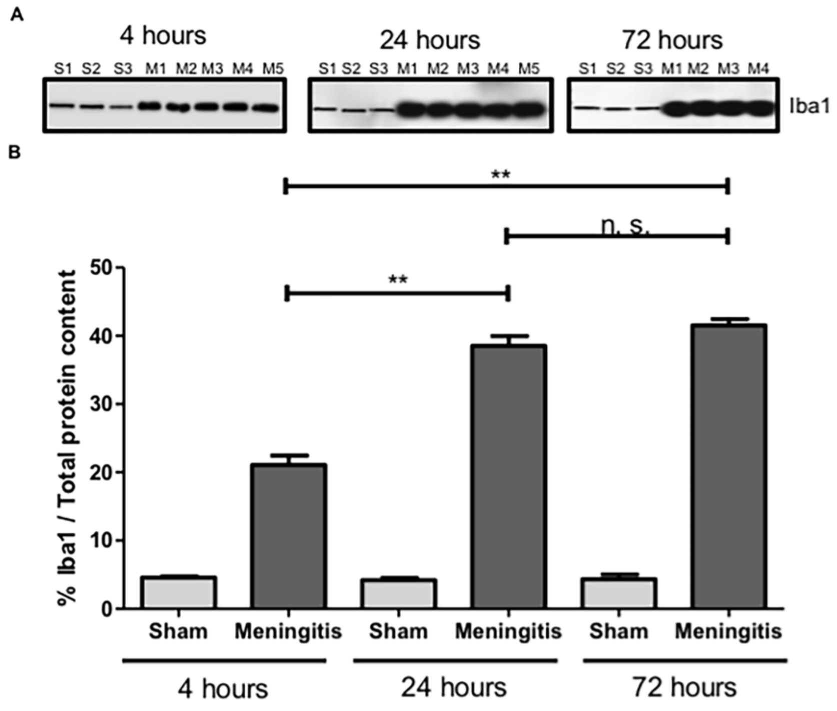

**Supplementary Figure S5. High-resolution immunofluorescence microscopy analysis of the astrocytic end feet detachment from the BBB vascular endothelium when the glymphatic system´s functionality is impaired.** High-resolution immunofluorescence microscopy analysis of BBB vascular endothelium in the cerebral cortex stained with Lycopersicon Esculentum tomato lectin DyLight 594 (red) and astrocytes stained with mouse anti-GFAP antibody combined with Alexa Fluor goat anti-mouse 488 (green). Brain tissue sections from sham rats clearly show astrocytes in close proximity around the brain vasculature in support of the BBB; in contrast, brain tissue from meningitis-affected rats showed a progressive detachment overtime of the astrocytic end feet from the vascular endothelium; the white double-arrows point towards the detaching astrocytic end feet; four brain tissue sections from three sham mice and five (four for the 72 hour-time-point) were analyzed per time point, ten images per each section were taken, the images displayed are representative of each group.

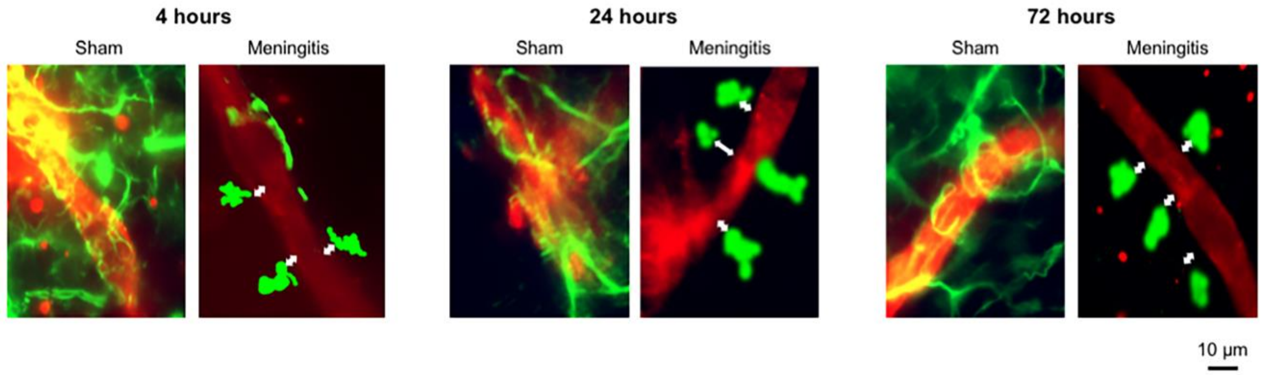

**Supplementary Table S1. Clinical score of the rats during experimental pneumococcal meningitis.** Body weight was on average 216.6 ± 4.5 g for the 4 hours' group; 218 ± 6 g for 24 hours, and 215.6 ± 3.8 g for 72 hours; in the column "CFU/mL in CSF", AV = average value for the Meningitis group in each time-point (all animals in the sham control group had negative bacterial cultures).

| Time | 1 = Coma | 2 = Does not turn upright when positioned on the back | 3 = Turns upright within 30 s | 4 = Minimal ambulatory activity, turns upright < 5 s | 5 = Normal | Eye secretion | Coat (piloerection) | Weight | CFU/mL in CSF (5 µl of CSF) |
|---|---|---|---|---|---|---|---|---|---|
| 4 hours | | | | | R1<br>R2<br>R3<br>R4<br>R5 | R1 | | R1 - 210 g<br>R2 - 222 g<br>R3 - 215 g<br>R4 - 219 g<br>R5 - 217 g | R1 $4.3x10^4$<br>R2 $1.5x10^5$<br>R3 $1.1x10^5$<br>R4 $6.6x10^4$<br>R5 $5.4x10^4$<br>**AV $8.5x10^4$** |
| 24 hours | | | | R1<br>R2<br>R3<br>R4<br>R5 | | | R1<br>R2<br>R3<br>R4 | R1 - 221 g<br>R2 - 215 g<br>R3 - 215 g<br>R4 - 227 g<br>R5 - 212 g | R1 $2.0x10^5$<br>R2 $7.4x10^5$<br>R3 $6.5x10^5$<br>R4 $6.8x10^5$<br>R5 $8.4\ x10^4$<br>**AV $4.7x10^5$** |
| 72 hours | | | | R1<br>R2<br>R3<br>R4<br>R5 | | | R1<br>R2<br>R3<br>R4<br>R5 | R1 - 210 g<br>R2 - 215 g<br>R3 - 215 g<br>R4 - 220 g<br>R5 - 218 g | R1 $6.9x10^5$<br>R2 $7.7x10^5$<br>R3 $8.5x10^5$<br>R4 $6.7x10^5$<br>R5 $1.8x10^6$<br>**AV $9.6x10^5$** |
